# Regulatory features determine the evolutionary fate of laterally acquired genes in plants

**DOI:** 10.1101/2025.08.22.671697

**Authors:** Catherine F Collins, Benjamin T Alston, Samuel GS Hibdige, Pauline Raimondeau, Emily Baker, Graciela Sotelo, Alexander S. T. Papadopulos, Pascal-Antoine Christin, Lara Pereira, Luke T Dunning

## Abstract

Lateral gene transfer (LGT) is widespread in eukaryotes, including in animals and plants where it can fuel adaptive evolution and innovation. However, the factors that influence the integration and long-term retention of transferred genes remain poorly understood. The pangenome of the grass *Alloteropsis* has a high turnover of laterally acquired genes, and here we combine expression, methylation and genomic data to identify factors promoting their long-term persistence. Most transferred genes appear to be degenerating, showing lower expression levels and/or greater sequence truncation compared to their vertically inherited homologs. These degenerating genes also show significantly higher levels of DNA methylation, potentially indicating transcriptional silencing. The likelihood of a transferred gene being retained will be influenced by how easily it can be expressed in the recipient genome. In *Alloteropsis*, putatively functional laterally acquired genes had expression levels significantly more similar to their donor xenolog than to their vertically inherited homolog. This pattern suggests that transferred genes may carry cis-regulatory elements encoded on the fragment of DNA that moves between species, facilitating their expression in the new genomic context. Evolutionary novelty may also increase the likelihood that selection retains a transferred gene. However, only a significant difference in expression level, not sequence divergence, between donor and recipient orthologs is associated with successful lateral gene transfer. Overall, our results show that most transferred genes degrade over time. However, those capable of regulating their own expression are more likely to persist and contribute to long-term evolutionary innovation.

**Significance Statement:** Lateral gene transfer (LGT) can introduce novel traits into plant genomes, yet most transferred genes are only transient residents and are degenerating, with reduced expression, truncation, and elevated DNA methylation. However, a minority persist and are more likely to resemble their donor counterparts in expression, suggesting co-transfer of cis-regulatory elements. These findings indicate that regulatory compatibility is key to their long-term survival.

## Introduction

Horizontal or lateral gene transfer (LGT) is the acquisition of genetic material outside of sexual reproduction^1^. The importance of this process for prokaryote evolution has long been established^2–4^, and over the last two decades it has become increasingly apparent that LGT can also play an important role in eukaryote evolution^5,6^. LGT effectively expands a species gene pool, enabling it to use genetic variation from other species^7^. This increased standing variation can accelerate adaptation^8^, and acquired genetic information has underpinned several evolutionary innovations in eukaryotes, including the colonisation of land by plants^9^, the silencing of host defences by parasites^10,11^, and the optimisation of core metabolism^12^. However, the specific genes transferred are random, and therefore the proportion of evolutionarily advantageous events is likely to be small. Indeed, in prokaryotes most transfers are neutral or deleterious, and rapidly lost in the recipient species through drift or selection^3,13,14^. While LGT is known to occur in a wide range of plants and animals, we are yet to test the factors that influence the successful integration and long-term retention of transferred genes.

The probability of successful LGT depends critically on whether the recipient can express the transferred gene and produce a functional protein^5^. LGT is particularly prevalent in parasitic plants^10,15–18^, where intimate physical contact with the host via the haustorium serves as a conduit for gene transfer. Despite the haustoria acting as a highway for mRNA movement between the species^19^, most transfers are surprisingly driven by the movement of genomic DNA^16^. This is thought to be because DNA fragments can also transfer intact promoters, increasing the likelihood that the gene remains functional in the recipient species^16^. In grasses^20^ and insects^21^, the number of successful transfers increases as phylogenetic distance decreases. This may also be due to ease-of-use, with more closely related species more likely to share regulatory mechanisms^20^. If the regulation of a gene is encoded on the transferred DNA fragment, then any advantageous regulatory feature from the donor may also be transferred. Therefore, genes which are predominantly influenced by cis-regulatory elements will be more easily expressed by the recipient and result in successful LGT.

The likelihood of retaining a laterally transferred gene will also increase if it adds functional novelty^5,22^, such as introducing a gene that the recipient lacks. For example, ferns acquired a chimeric neochrome photoreceptor that was unique to hornworts, enabling them to successfully grow in low-light conditions^23^. However, not all transfers involve novel acquisitions. In grasses, nearly 80% of transferred genes co-exist with a vertically inherited homolog in the recipient’s genome^20^. In these cases, functional novelty may arise from genetic variation that accumulated since the donor and recipient lineages diverged. These differences can accumulate neutrally over time through genetic drift, and when the gene is transferred into a new genomic environment, it can enhance standing genetic variation that can later fuel adaptation^24^. Alternatively, the donor ortholog may have undergone episodic positive selection. For example, phosphoenolpyruvate carboxylase (PEPC) is a key enzyme in C_4_ photosynthesis, an adaptive trait that increases carbon fixation in hot and high light environments^25^. Three different genes encoding this enzyme were laterally acquired by separate populations of the grass *Alloteropsis semialata*^12,26^. These genes had been optimised for C_4_ metabolism in older C_4_ grass lineages through positive selection^26^, and their recurrent acquisition accelerated metabolic adaptation in the recipient^27^. While there is some anecdotal evidence of its importance in eukaryotes, it has yet to be formally tested whether the likelihood of retaining a laterally acquired gene is correlated with the functional novelty it provides to the recipient.

LGT appears to be particularly widespread in the grass family^12,20,28–35^, making it a useful system to determine the factors that influence the integration and retention of laterally acquired genes. An in-depth phylogenetic analysis of grass-to-grass transfers in the *Alloteropsis* pangenome identified 168 unique laterally acquired genes in five reference genomes (Table 1), revealing a high turnover with continual gene gain and loss^35^. The distribution of the laterally acquired genes was then determined across the lineage using additional whole genome re-sequencing datasets, indicating that only a small proportion are old acquisitions retained over time, and most are relatively recent and geographically restricted^35^. Their apparent transience is further supported by the rate at which they are lost from the recipient species, which is more than 500 times higher than that of vertically inherited genes^35^.

**Table 1:** The number of laterally acquired genes identified in *Alloteropsis*. The laterally acquired genes were identified by Raimondeau, et al., (2023) using both read mapping and phylogenetic approaches, and in this study we restricted our analysis to only include those identified by both methods, were present in the original genome annotation, and were not 100% identical to another sequence. For each accession the number of unique laterally acquired genes is shown, and the number of genes this corresponds to in the genome annotation is given in parenthesis (this counts duplicates that could have arisen prior or post lateral transfer separately). The total row indicates the number of unique laterally acquired genes across the *Alloteropsis* lineage, meaning that those shared by multiple reference genomes are only counted once. The number in the brackets on this row counts the laterally acquired genes in all five reference genomes, and their duplicates, separately.

| Species | Accession | Location | Raimondeau et at. (2023) |  |  |
| --- | --- | --- | --- | --- | --- |
|  |  |  | Read mapping | phylogenetics | Used in this study |
| *A. semialata* | AUS1 | Australia | 50 | 50 [53] | 50 [52] |
|  | RSA5-03 | South Africa | 33 | 29 [31] | 26 [28] |
|  | TAN1-04B | Tanzania | 45 | 43 [51] | 43 [51] |
|  | ZAM1505-10 | Zambia | 100 | 95 [125] | 94 [124] |
| *A. angusta* | UGA4 | Uganda | 32 | 32 [33] | 26 [27] |
| Total |  |  | 168 | 168 [293] | 167 [282] |

Here, we expand on this previous study in *Alloteropsis* by comparing the patterns of expression and sequence evolution of the laterally acquired genes identified in the *Alloteropsis* pangenome with the xenologs in the donor species and vertically inherited homologs in the recipient. Using these data we specifically test the following hypotheses: [1] laterally acquired genes are generally expressed at lower levels, more truncated and more heavily methylated that the corresponding homolog in the recipient species, consistent with the expectation that most are not adaptive and are undergoing functional degradation. [2] For the minority of laterally acquired genes that remain functional, their expression patterns will more closely resemble those of the donor xenolog than the recipient homolog, suggesting that cis-regulatatory elements are important for their retention. Finally, [3] functional laterally acquired genes will correspond to orthologs with greater sequence and expression divergence between donor and recipient than those that degenerate post-acquisition, suggesting evolutionary novelty increases their chances of long-term retention.

## Results

### Laterally acquired genes are expressed at relatively low levels

We generated 55 Illumina RNA-Seq data sets for the same five *Alloteropsis* accessions that have reference genomes and were used by Raimondeau *et al*.^35^ to identify laterally acquired genes (Table 1). For each accession, we sequenced up to three replicated samples from four tissues (leaf base, leaf tip, leaf sheath and root) (Table S1). To infer gene expression levels (transcripts per million; TPM) we used a reference-based approach, mapping each accession to its respective reference genomes. Out of the 282 individual laterally acquired genes identified by both approaches in Raimondeau et al.^35^, and present in the original genome annotations (Table 1), 208 (73.8%) were expressed (≥ 0.5 TPM) in at least one of the 55 RNA-Seq datasets, with a similar proportion expressed in each of the five reference genomes (mean = 77.5%; SD = 8.3%; Figure 1; Table S2). Most laterally acquired genes co-exist in the genome with their vertically inherited homolog (per accession mean = 72.1%, SD = 10.7%), with a significantly higher proportion (p = 0.008; Wilcoxon rank sum test) of recent duplication in the latter (laterally acquired genes: per accession mean = 10.3%; SD = 9.1%; vertical homolog: per accession mean = 31.9%; SD = 9.1%). The presence of a vertical homolog is not associated with the expression of laterally acquired genes (p = 0.830; Pearson Chi-squared test).

**Figure 1:**
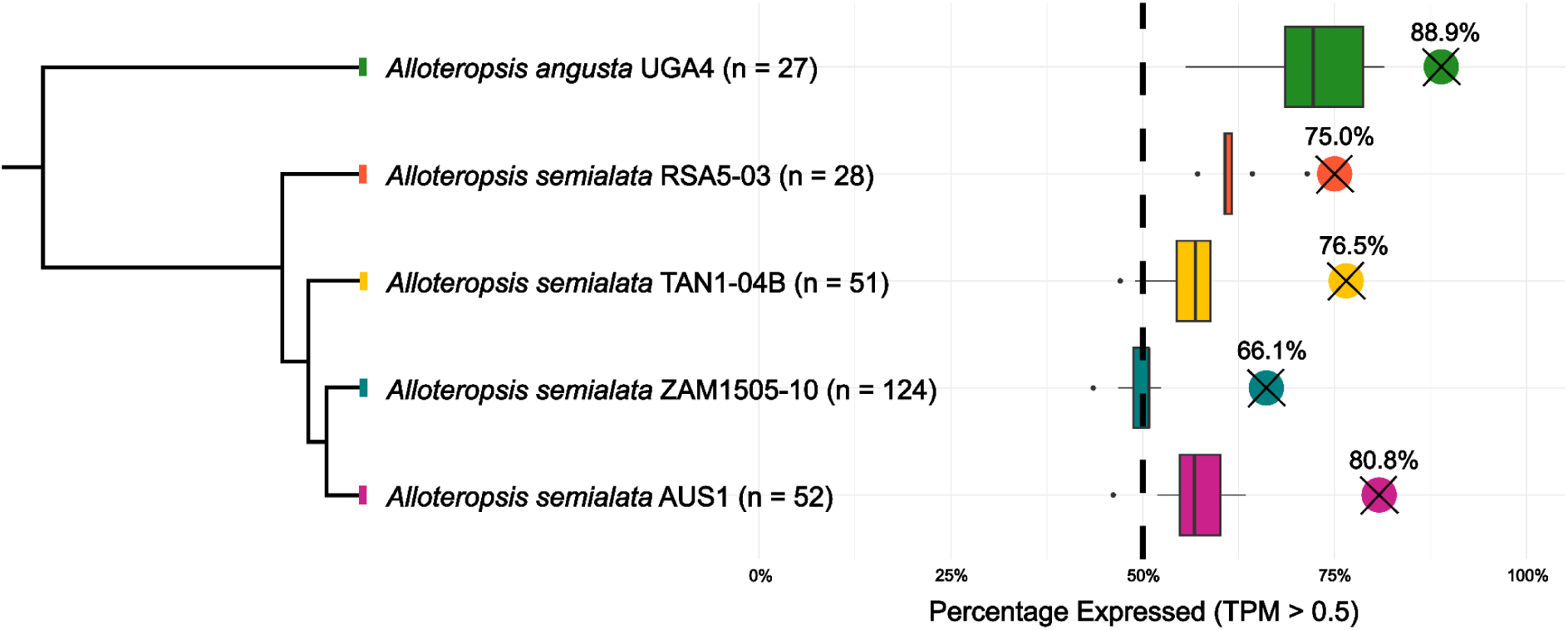
Expression of laterally acquired genes in *Alloteropsis*. A phylogenetic tree adapted from Bianconi et al.^67^ shows the evolutionary relationships of the *Alloteropsis* accessions, with the number of laterally acquired genes in each genome indicated (n). The dots with crosses show the percentage of genes where at least one replicate passes the expression threshold (TPM > 0.5) in any tissue. Boxplots show the median and interquartile range of the number of laterally acquired genes expressed in each individual RNA-seq dataset.

To compare the overall expression of laterally acquired genes and their vertically inherited homologs, we first calculated the log-TPM difference between them independently for all RNA-seq datasets from the different accessions and tissues (Figure 2a). The log-TPM differences were then combined into a single linear mixed-effects model (LMM) that included accession and orthology to account for non-independence (a similar approach was taken for all linear models). Overall, laterally acquired genes are expressed at significantly lower levels compared to vertically inherited homologs (p = 1.87 × 10^-14^; Figure 2a). Using the same approach with an additional 19 RNA-seq datasets from the donor species (Table S1), we also showed that the laterally acquired genes are generally expressed at significantly lower levels (LMM; p < 2 x 10^-16^; Figure 2b) than their xenolog in the donor. For the donor analysis, we used *Themeda triandra* as a proxy for transfers from Andropogoneae species (n = 42 genes in the 5 genomes), and *Setaria italica* for transfers from Cenchrinae species (n = 69 in the 5 genomes).

**Figure 2:**
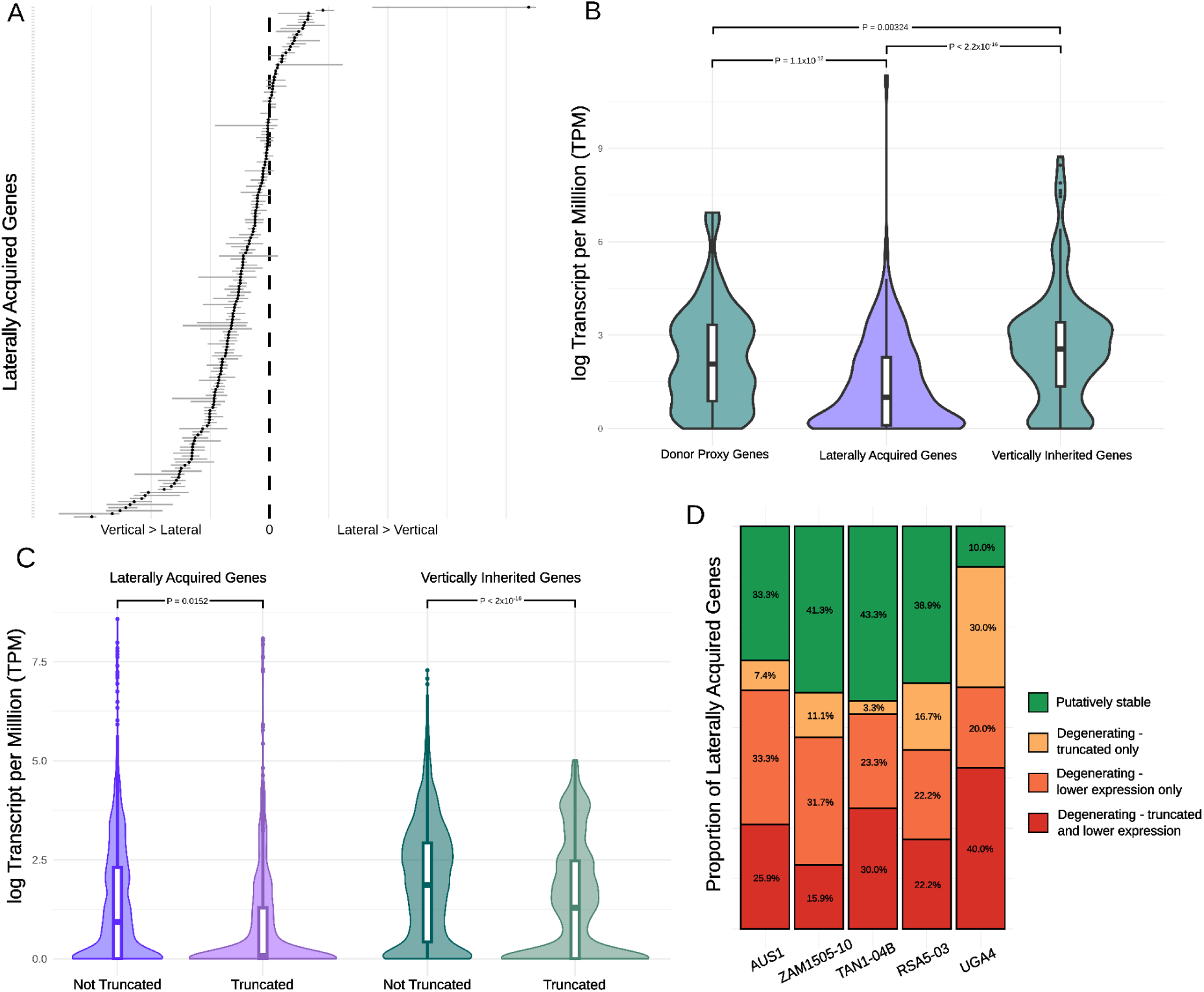
Expression patterns of laterally acquired genes. A) Comparison of expression levels between laterally acquired and vertically inherited homologs, all tissues combined. For each gene pair (n = 169) in each replicate, the log-transformed expression of the vertically inherited gene was subtracted from that of its corresponding laterally acquired homolog. Each dot indicates the median for the laterally acquired gene pair’s replicates (paralog expression summed), the horizontal indicates interquartile range. The vertical dotted line corresponds to a difference of zero between lateral and vertical expression. B) Comparison of log-transformed expression levels between laterally acquired, vertically inherited and donor xenologs, with significant differences indicated. Laterally acquired genes show a mean 59.7% decrease in expression compared to the vertically inherited homologs, and 40.6% decrease compared to the donor xenologs. C) Comparison in log-transformed expression levels between truncated and non-truncated genes, for laterally acquired and vertically inherited homologs. D) Laterally acquired genes were divided into four categories based on whether they (i) had significantly lower expression levels than the vertically inherited and donor proxy gene and (ii) were classified as truncated (< 70% sequence length in comparison to nearest orthologs).

There were 107 laterally acquired genes (range of 10-63 per accession) for which both the vertically inherited homolog and donor xenolog were present and expressed (TPM > 0.5; Table S2). When the laterally acquired genes were compared to both of these combining all tissues, over half (per accession mean = 53.8%; SD = 6.1%) were expressed at a significantly lower level (adjusted p < 0.05; Wilcoxon rank sum test; Table S3). Those with significantly reduced expression were classified as “degenerating”, with either the transfer itself inhibiting activity, or the gene being actively downregulated post acquisition. The remaining genes were classified as “putatively stable”. In total, 69.6% of the laterally acquired genes shared between accessions were placed in the same category (Table S3).

### Laterally acquired genes have higher gene body methylation

Epigenetic silencing is one mechanism by which transgenes are rapidly silenced in genetically modified plants^36^, and laterally acquired genes may be subjected to similar processes. To investigate this, we generated bisulfite sequencing data from leaf tissue for the four *A. semialata* accessions, and calculated the proportion of methylated cytosines (CpG) for the gene body, exons, and 1kb up- and down-stream. To test for a difference in the proportion of CpG methylation between laterally acquired genes and their vertically inherited homologs, we again used a LMM. Laterally acquired genes had significantly higher CpG methylation across the gene body (p = 0.001), exons (p = 0.004), and 1 kb down-stream (p = 0.026; Figure 3a). This difference was most pronounced across the gene body, with a mean of 2.7% increase in CpG methylated sites (Figure 3a).

**Figure 3:**
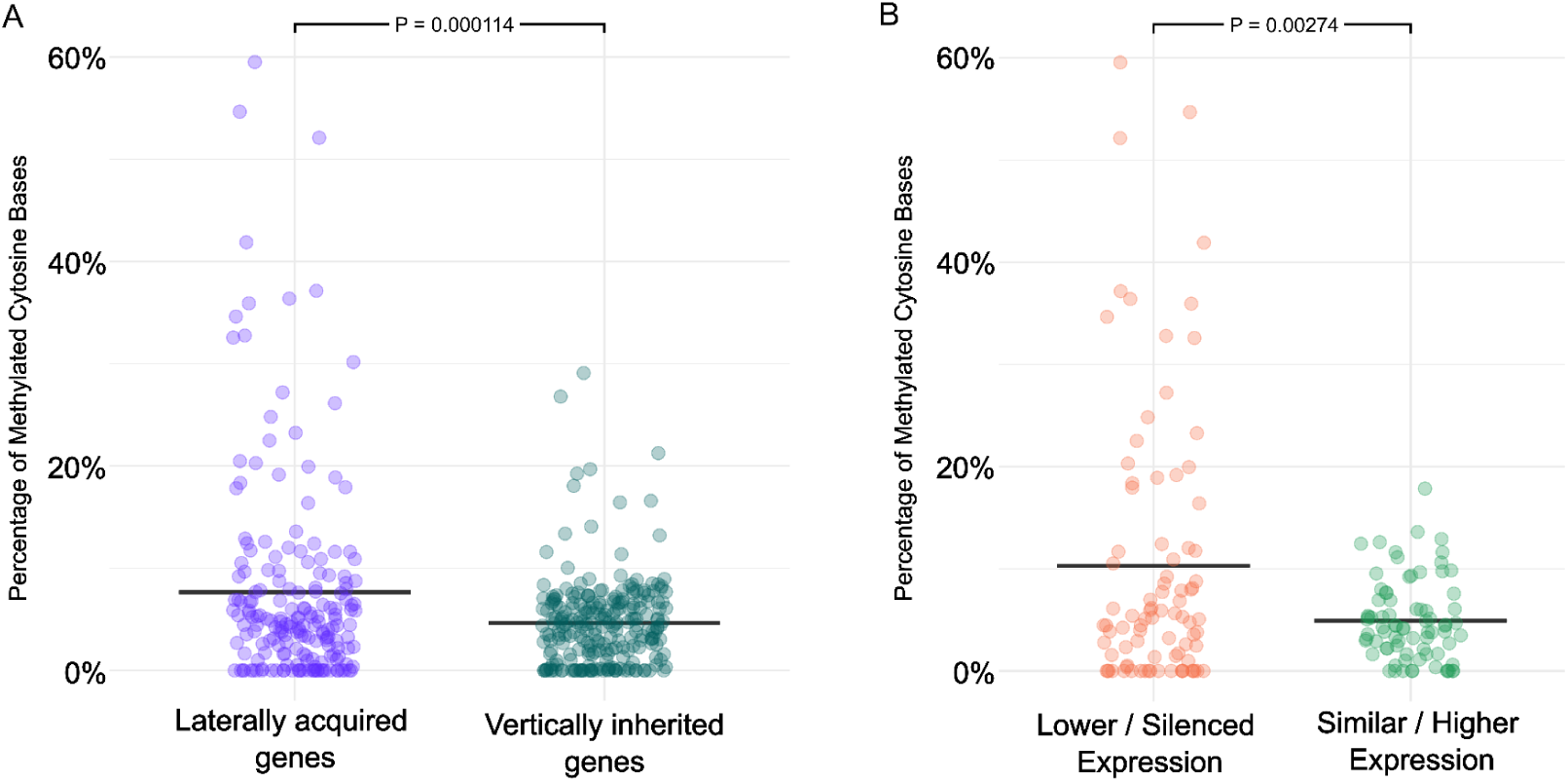
Methylation of laterally acquired genes. Comparison of proportion of methylated cytosine sites across the full gene region between a) laterally acquired and vertically inherited genes and b) laterally acquired genes with silenced or lower expression than their corresponding vertically inherited and donor proxy genes, and those with similar or higher expression. Horizontal bars indicate the mean.

We repeated the LMM analysis to compare CpG methylation between the degenerating and putatively stable laterally acquired genes. Those that are classed as degenerating had a significantly higher proportion of CpG sites in the gene body (p = 0.003; Figure 3b). This pattern was also observed on the exons (p = 0.034).

### Laterally acquired genes are frequently truncated

The coding sequences of laterally acquired genes and their vertically inherited homologs in the *Alloteropsis* reference genomes were compared with those of other model grasses to assess whether either were truncated. They were classed as truncated if: (a) the gene was less than 70% of the length of all other orthologs, (b) the annotated sequence did not begin with a start codon, or (c) the annotated sequence did not end with a stop codon. Across all 5 genomes, 33.3% of the laterally acquired genes were classed as truncated compared to 16.2% of their vertically inherited homologs. Even though there were 10 truncated vertically inherited homologs with full length laterally acquired genes, the increase in the overall proportion of full length laterally acquired genes with truncated vertically inherited homologs was not significant (p = 0.607). A generalised linear mixed model (GLMM) indicated that vertically inherited homologs have significantly lower odds of being truncated than laterally acquired genes (p = 4.44 x 10^-4^). A separate LMM also showed that truncated genes had significantly lower expression levels than non-truncated genes (p = 3.43 x 10^-16^; Figure 2c), n.b. The TPM expression measure accounts for gene length. Finally, degenerating genes were significantly more likely to be truncated than those that are putatively stable (GLMM p = 0.012). Given structural decay suggests functional loss, we reclassified truncated genes previously considered putatively stable as degenerating (if paralogs were present all copies had to be truncated). This adjustment further demonstrates that a substantial majority of laterally acquired genes are degenerating (mean = 66.6%, SD = 13.6%; Figure 2d).

### Functional laterally acquired genes have donor-like expression levels

We conducted a more detailed analysis on the expression pattern of the 56 laterally acquired genes that appear to be putatively stable in *Alloteropsis* to test if their expression pattern was more similar to the vertically inherited homolog or to the xenolog in the donor (Figure 4a). The 56 laterally acquired genes represent 45 unique genes when accounting for orthologs present in more than one *Alloteropsis* accession. To test for a significant difference between the laterally acquired genes and their vertically inherited homolog/donor xenolog we used Kruskal-Wallis tests (FDR-corrected), followed by Dunn’s post hoc test for pairwise differences using the FSA package^37^ in R. Based on these results, the laterally acquired genes were then placed into one of five different categories: (1) ‘No difference’ - no significant difference among the three groups; (2) ‘Higher’ - expression significantly higher than both donor and vertical xenologs, (3) ‘Intermediate’ - expression significantly different from both, but intermediate, and either (4) ‘recipient-like’ or (5) ‘donor-like’ if significantly different from only one but not the other. We then used binomial tests to look for significant differences in the number of genes assigned to each category.

**Figure 4:**
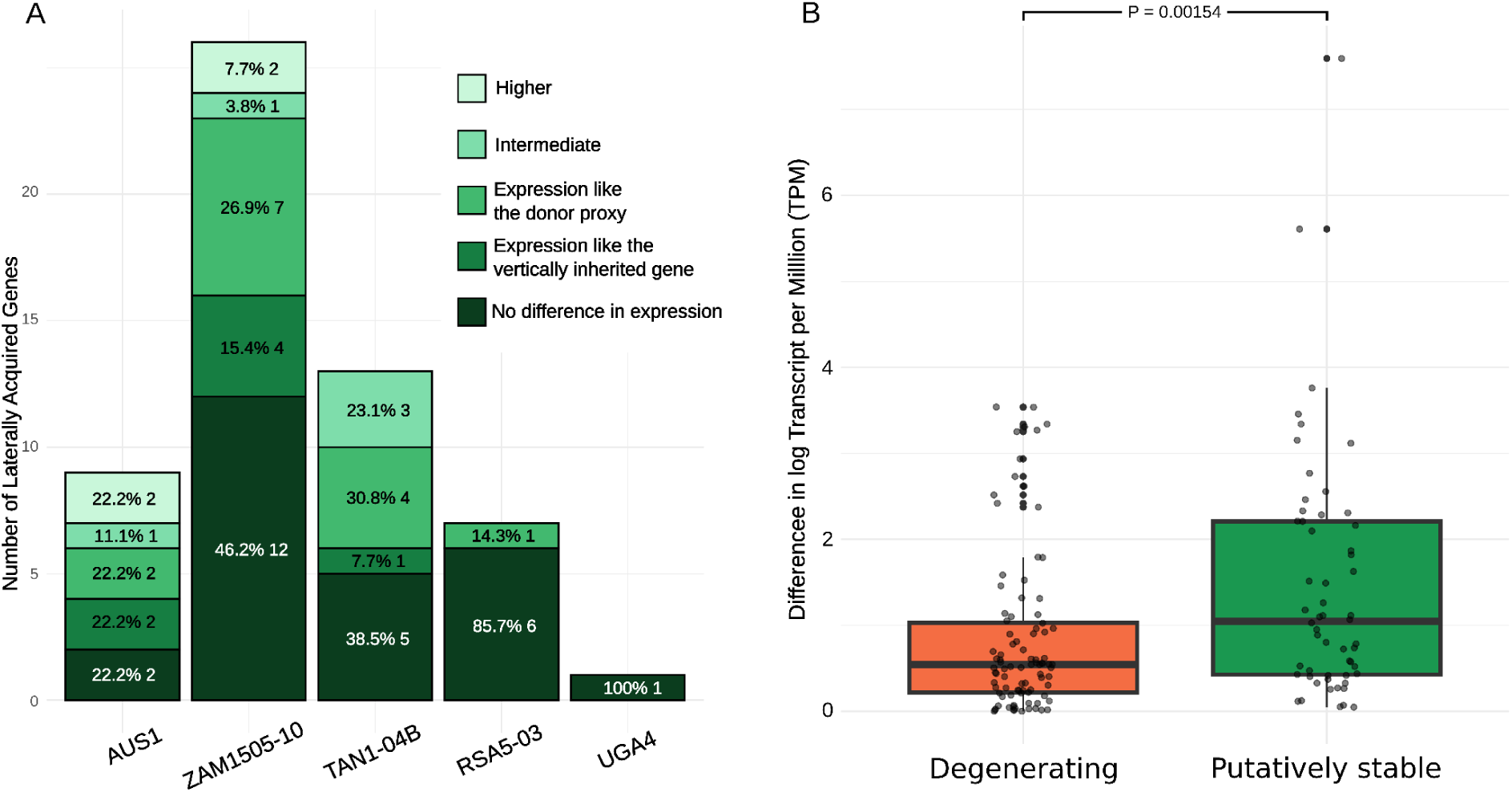
Expression patterns using vertically inherited and donor proxy genes as a comparison. A) Expression of laterally acquired genes classified as ‘putatively stable’, i.e. gene expression of the gene is not significantly lower than both the vertically inherited homolog and donor xenolog, and it has not been classified as truncated. Categories refer to the relationship between the laterally acquired gene and its corresponding vertically inherited homolog and donor xenologs. Higher: laterally acquired gene significantly higher in expression than the corresponding homologous genes; intermediate: laterally acquired gene intermediate in expression between the homologous genes. B) Difference in log transformed expression levels between corresponding vertically inherited and donor proxy genes for laterally acquired genes classified as Degenerating or ‘Putatively stable’. Comparison of log transformed expression counts between vertically inherited and donor proxy genes corresponding to laterally acquired genes classified as ‘Degenerating’ or ‘Putatively stable’. P value from Wilcoxon Rank Sum test is shown above boxplots.

A majority of putatively stable laterally acquired genes showed no difference in expression to the vertically inherited gene or to the donor xenologs (p = 7.58 x 10^-6^, exact binomial test). The ‘no difference’ category is not informative as to whether the laterally acquired gene expression level is dictated by cis-acting regulatory elements linked to the laterally acquired gene DNA fragment, or if its expression level is now governed by the recipient’s own regulatory machinery. We therefore repeated the analysis without this classification, and out of the remaining four categories, putatively stable genes where expression levels were more similar to the donor proxy were significantly over-represented (p = 0.008, exact binomial test). This pattern is largely driven by three out of the five accessions (TAN1-04B, UGA4 and ZAM1505-10) (Figure 4a).

### Evidence of post-transfer expression modification

Three *Alloteropsis semialata* accessions possess a laterally acquired phosphoenolpyruvate carboxykinase (PCK) gene, which forms part of the C_4_ photosynthetic pathway^12^. In the exclusively C_4_ accessions ZAM1505-10 and AUS1, expression of this laterally acquired PCK in the leaf and sheath tissues is comparable to the donor PCK (p = 0.232; p = 0.885). However, it is expressed significantly lower than the donor and vertical PCK in the roots (p = 4.76 x 10^-6^; p = 0.004). In these accessions, the vertically inherited PCK has lower expression in leaf and sheath (p = 5.34 x 10^6^; p = 0.002), but is expressed at similar levels to the donor PCK in the root (p = 0.262; p = 0.138. This suggests that the transfer of PCK has driven subfunctionalization, with the laterally acquired PCK now only responsible for C_4_ photosynthesis, and the vertically inherited PCK restricted to a ubiquitous housekeeping role. Contrasting results were observed in TAN1-04B, a C_3_+C_4_ intermediate, in which only part of its carbon fixation is through the C_4_ pathway that requires high PCK expression. In this case, the laterally acquired PCK gene shows similar expression to the vertically inherited copy (p = 0.841), both being expressed significantly lower than the donor PCK (p = 9.71 x 10^5^).

### Stable transfers have donor and recipient orthologs with increased expression divergence

We calculated the absolute difference in mean expression (logTPM) between vertically inherited and donor orthologs for each laterally acquired gene (Table S4). The differences were significantly larger for orthologs associated with ‘putatively stable’ (mean = 1.46; SD = 1.43) rather than ‘degenerating’ (mean = 0.844; SD = 0.944) lateral acquired genes (Wilcoxon rank-sum test: p = 0.002; Figure 4b). We also fitted a LMM to test the interaction between gene status (degenerating vs putatively stable) and gene type (vertically inherited vs donor) on expression levels, including accession and orthology to account for non-independence. The interaction was highly significant (p = 9.52 x 10^-12^), confirming that difference in gene expression between the vertically inherited and donor homolog is significantly greater in putatively stable laterally acquired genes than in degenerating ones. For putatively stable genes, vertical gene expression is generally higher than donor gene expression (Table S4).

### Stable transfers do not have increased nucleotide divergence between donor and recipient orthologs

We calculated the pairwise neutral genetic divergence (the rate of synonymous substitutions [dS]) between vertically inherited and donor orthologs for each laterally acquired gene (Table S5). There was no significant difference between orthologs associated with ‘putatively stable’ (mean = 0.447; SD = 0.379) or ‘degenerating’ (mean = 0.474; SD = 0.568) lateral acquired genes (Wilcoxon rank-sum test: p = 0.837; Figure S3). There was also no difference between the rate of non-synonymous substitutions (dN, Wilcoxon rank-sum test: p = 0.658) or the overall selection pressure (dN/dS, Wilcoxon rank-sum test: p = 0.645). LMM’s also showed no significant interaction between gene status (degenerating vs putatively stable) and gene type (vertically inherited vs donor) for dN (p = 0.641), dS (p = 0.575) or dN/dS (p = 0.89).

### Laterally acquired genes show increased relaxed selection compared to the vertically inherited homolog

Using branch and branch-site models in CodeML^38^, we calculated the dN/dS ratio (ω) for laterally acquired and vertically inherited homologs since they coalesced in the gene tree (Table S5). For the laterally acquired gene, the ω value reflects its history of selection in the donor lineage and the time since it was transferred into the recipient genome. The branch model identified significant shifts in ω for 39/116 (14 decreased ω, 25 increased ω) laterally acquired genes (P<0.05; Figure 5a), with similar proportions for those classified as degenerating (10.8% decreased ω, 24.1% increased ω) and functional (11.5% decreased ω, 19.2% increased ω). In comparison, only 14/116 vertically inherited homologs showed significant shifts in ω compared to the background tree (p < 0.05; 10 decreased ω, 4 increased ω; Figure 5a). Subsequent branch-site analysis of the genes showing relaxed selection (increased ω) identified significant positive selection in 6/25 (3 degenerating, 2 stable, 1 with no identified donor proxy gene) laterally acquired genes, and 2/4 of the vertically inherited genes (p < 0.05; Figure 5b).

**Figure 5:**
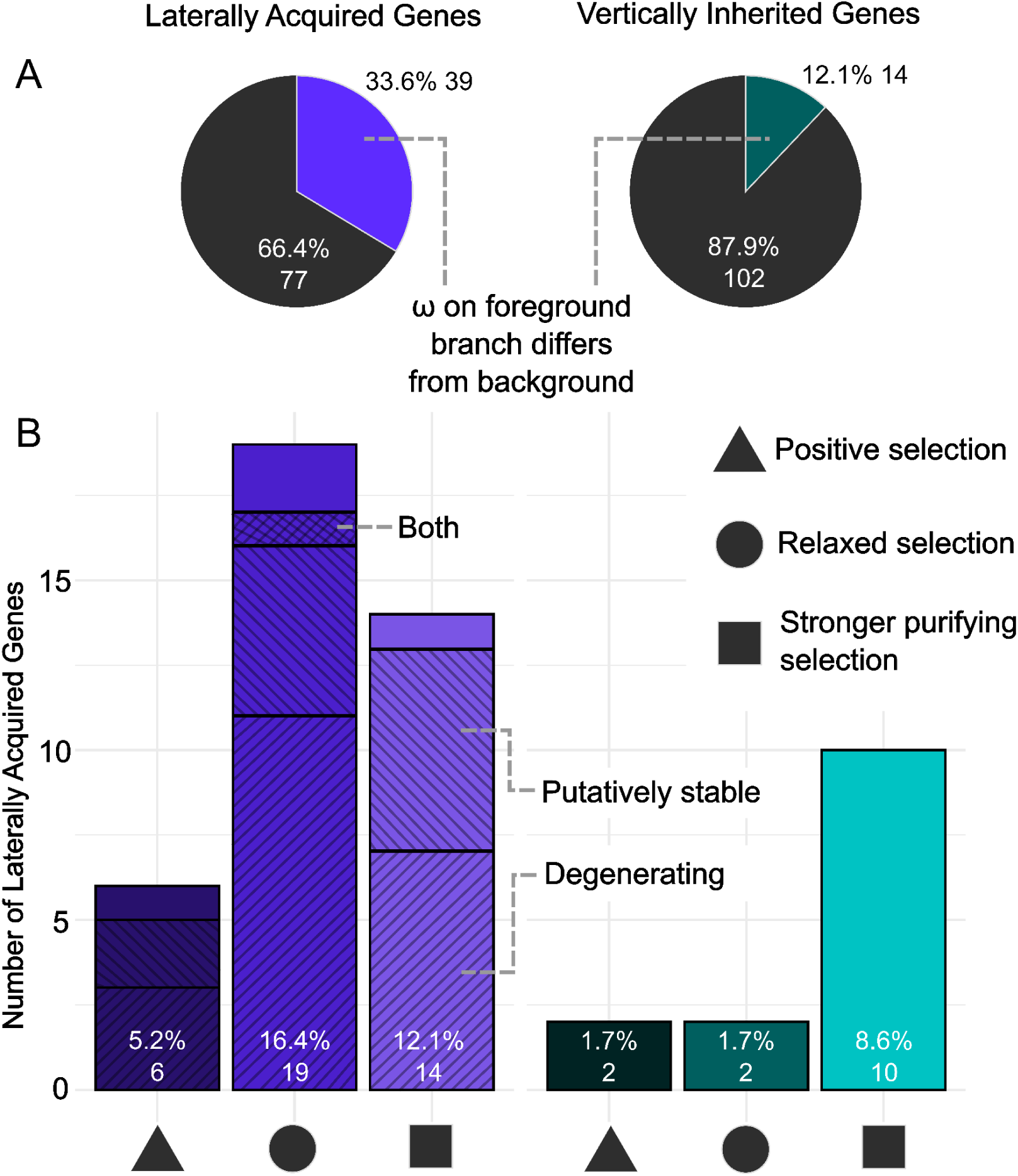
Selection patterns in laterally acquired and vertically inherited genes. A) Colored segments indicate genes where the foreground branch experienced significantly different selection pressures compared to background branches, as determined by the CodeML branch model (p < 0.05). B) Genes significantly different were further categorised as: stronger purifying selection, where the dN/dS ratio (ω) for the gene is lower than the background ω; positive selection, where ω for the gene is higher than the background ω and evidence for sites under positive selection has been found using the PAML CodeML branch site model (p < 0.05); and relaxed selection, where ω for the gene is greater than the background ω but not significant in the branch site model. Percentages represent the proportion of overall genes (n = 116) in each category. For laterally acquired genes, hashing indicates ‘degenerating’ and ‘putatively stable’ categorisation. ‘Both’ indicates where categorisation differed across accessions. No hash indicates genes where a donor copy was not present.

Overall, laterally acquired genes were significantly more likely to show relaxed constraint than vertically inherited genes (Fisher’s exact test, p = 1.12 x 10^-4^), although there was no difference between those classified as degenerating or putatively functional (Fisher’s exact test, p = 0.503). It should be noted that these results do not include the three laterally acquired PEPC genes in *Alloteropsis* that were acquired from Andropogoneae, Cenchrinae and Melinidinae species, and have been shown to contain positively selected amino acid residues^12^. This exclusion was because in Raimondeau et al.^35^ the genes either did not pass the stringent phylogenetic filters used to identify LGT (Cenchrinae and Melinidinae PEPC in ZAM1505-10), or the inferred gene tree lacked a vertically inherited copy in the same accession for comparison (Andropogoneae PEPC in AUS1).

### Time since acquisition is not correlated with expression level

Molecular dates for the timing of each acquisition were obtained from Raimondeau et al.^35^ (Table S6). Overall, laterally acquired genes tend to be young, with 86 genes (54.4%) less than 2 million years old (Ma), 36 genes (22.8%) in the 2–4 Ma range, 32 genes (20.3%) in the 4–6 Ma range, and only 14 genes (8.9%) in the ‘more than 6 Ma’ category. Although overall there is a correlation between the time since acquisition and expression level (LM; p = 3.56 x 10^-5^; R^2^ = 0.005), this relationship is not significant when accounting for non-independence in the data (orthology and accession). At the accession level, significant time–expression level relationships were detected in AUS1 (LM p < 2 × 10⁻¹⁶; LMER p = 0.00463), TAN1-04B (LM p = 0.000753), ZAM1505-10 (LM p = 3.64 × 10⁻⁶), and USA4 (LM p = 0.000259; Figure S3). KWT showed no significant associations in either model. When age was compared between ‘Degenerating’ and ‘Putatively stable’, no significant difference was found (p = 0.1964; Wilcoxon Rank Sum Test). At the accession level, only ZAM1505-10 found a significant difference in age between these categories (p = 0.0172); all other accessions showed no significant difference (Figure S3).

## Discussion

### Most laterally gene transfers are unsuccessful

Most laterally acquired genes are expressed, however, a majority show significantly reduced expression compared to both their vertically inherited homolog and donor xenolog. This suggests that either the transfer of the gene itself has inhibited activity, or the gene has been downregulated following acquisition. The transferred genes also have a significantly higher sequence truncation rate (> 2-fold) in the genome compared to their vertically inherited counterparts. We collectively termed the relatively lowly expressed and/or truncated genes as “degenerating”, as their reduced expression and structural decay suggest functional loss. Degenerating genes represented a substantial proportion of the lateral gene content across the five genomes (mean = 68.3%, SD = 13.0%; Figure 2d). The abolition of gene expression and nonsense mutations that generate truncated proteins are both processes that lead to pseudogenisation and ultimately the loss of a gene from a genome^39^. Previous findings in *Alloteropsi*s have shown that laterally acquired genes are lost at a rate 500 times higher than vertically inherited loci^35^. Most laterally acquired genes in *Alloteropsis* appear to be non-adaptive and are only transient components of the recipient’s genome, mirroring patterns previously seen in prokaryotes^3,13,14^. This means the genes detected likely represent only the briefly visible fraction of a much more extensive and ongoing process of lateral gene transfer^5^, with only a small subset occasionally retained through positive selection^24^.

Epigenetic silencing pathways can target and rapidly suppress transgenic DNA^40^, and a similar mechanism may act to reduce the expression of laterally acquired genes, which are functionally analogous to transgenes. In *Alloteropsis*, the laterally acquired genes have significantly higher gene body methylation than their vertically inherited homologs. Furthermore, degenerating transferred genes have significantly more gene body methylation than those that remain transcriptionally active (Figure 3). However, there is no significant difference in the number of methylated sites in the 1 kb upstream promoter region, a region typically associated with transcriptional suppression^41^. Given that these transfers happened many thousand or even millions of years ago, the promoter methylation may have been rendered obsolete in this time. Whilst the function of gene body methylation is debated^42^, methylation levels are generally associated with chromatin accessibility^43^, and elevated gene body methylation may reflect chromatin remodelling inhibiting the activity of the transferred genes. This may reduce the expression and any potential deleterious effect the gene has, rendering its persistence in the genome subject to neutral evolutionary forces.

### Successful transfers encode their own regulation

The putatively functional laterally acquired genes exhibit expression patterns that are more often similar to the donor xenolog than to the vertically inherited homolog (p = 0.008, exact binomial test; Figure 4a). The genes themselves are transferred as part of large DNA fragments^32^, and this result implies that successful transfers are more likely to be transcriptional units that carry linked cis-regulatory elements, such as promoters. This process mirrors successful transgenesis, in which the promoter sequences are critical for regulating the spatial and temporal expression of the transgene^44^. Genes that heavily depend on trans-regulatatory mechanisms are less likely to be functionally retained after transfer, as they will become decoupled from the necessary regulatory landscape during the process. Cis-regulatatory differences are also more commonly responsible for adaptive evolution^45^.

Whilst initial ‘*plug in and play*’ may have an important role in successful LGT, we do see evidence of post-acquisition modification of gene expression patterns. PCK is present in plants where it is ubiquitously expressed in all tissues and plays a role in gluconeogenesis. In some species, PCK is also co-opted for the C_4_ photosynthetic cycle, meaning that it becomes very highly expressed in leaf tissue. In the C_4_ *Alloteropsis,* the lateral acquisition of an additional gene encoding PCK has enabled subfunctionalisation. The vertically inherited copy retains its ancestral gluconeogenesis function (moderately expressed in leaf and root tissue), whereas the laterally acquired copy is specialised C_4_ photosynthesis (highly expressed in leaves, lowly expressed in roots) (SI Appendix, Figure S1). This post-acquisition specialisation demonstrates that both the genomic and regulatory contexts into which a gene is inserted play a critical role in shaping its evolutionary fate. Furthermore, the likelihood of retaining a transferred gene may increase if the vertically inherited homolog was under balancing selection for multiple functions prior to the initial transfer event.

### Functional novelty of gene expression profiles

The likelihood of successful lateral gene transfer increases when it introduces functional novelty that is maintained by selection^5^. Although the underlying mechanisms differ, this also applies to adaptive introgression, such as the spread of insecticide resistant genes in mosquitoes^46^. Functional novelty can arise through divergence in the coding sequence or how the gene is expressed. Here, we find that functional lateral gene transfers are more common for genes exhibiting greater expression divergence between donor and recipient orthologs (Figure 4b). The acquisition of genes with novel expression patterns can be adaptive, for example the introgression of a gene with a cis-regulatory mutation has been associated with adaptive shifts in flower colour in the monkeyflower *Mimulus aurantiacus*^47^.

In contrast, functional lateral gene transfers are not enriched for genes with elevated coding sequence divergence between donor and recipient orthologs (Figure S2). The relative contribution of protein-coding versus regulatory change to adaptive evolution has been debated, but data from Murid rodents suggests while adaptive noncoding regulatory changes may be more frequent, substitutions in proteins may have the largest effect on phenotypic evolution^48^. This may explain the overall enrichment of functional genes with expression, but not coding, divergence, as the increased frequency of the former means the recipient is more likely to acquire genes with adaptive regulatory variation that can be selectively maintained.

Selection for transferred genes with adaptive protein coding divergence is still likely to be important in their retention. For example, the repeated acquisition of genes encoding PEPC by *Alloteropsis*, which were subject to strong positive selection in the donor lineages^12,26^. Our results did show that laterally acquired genes were more likely to be under relaxed selection than vertically inherited genes, possibly reflecting a shift to relaxed selection in the recipient as they are likely either deleterious or neutral. However, we did detect positive selection in 3-fold more laterally acquired genes compared to vertical inherited homologs (Figure 5), but it should be noted that the absolute numbers are extremely low and there was no significant difference between those that appeared to be functional versus degenerating.

### Time since transfer

Based on our finding that most laterally acquired genes are degenerating, we would expect their expression levels to decay over time, with only those selectively retained remaining expressed in the long-term. However, overall we found no significant correlation between time since transfer and either expression levels or the classification of genes as either degenerating or putatively stable (Figure S3), although it should be noted that there were significant associations at the individual level for some accessions (Figure S3). Several factors could explain this pattern. Laterally acquired genes are typically transferred as part of large genomic fragments^32^, potentially allowing neutral or slightly deleterious genes to persist through hitchhiking with beneficial ones^24^. Similarly, if inserted into regions of the genome with a lower recombination rate it may take longer for them to be silenced and purged. In both instances, selection will take even longer to act if the laterally acquired gene is weakly deleterious or even selectively neutral. Finally, fluctuating selection in changing environments can maintain genetic variation, with genes initially retained for their adaptive potential but later becoming non-functional^49^. Similarly, it has been shown that one hitchhiking gene in *A. semialata* had a delayed selective impact, contributing to standing variation that became advantageous in a different geographic context^24^.

## Conclusion

Lateral gene transfer (LGT) in *Alloteropsis* is frequent but largely transient, with most acquired genes showing signs of degradation through reduced expression and structural decay. However, a subset of acquired genes remain functional, particularly those with donor-like expression. This indicates that the presence of cis-regulatory elements on the fragments of DNA moving between species may increase the chances of their retention. The functional laterally acquired genes also have donor and recipient orthologs with greater expression divergence, potentially indicating selection for regulatory novelty. The lack of a clear relationship between gene age and degeneration suggests that factors such as genomic context, fluctuating selection, and hitchhiking may mediate their long-term retention. Overall, while most laterally acquired genes are short-lived in the recipient species, those that persist can contribute to regulatory innovation and adaptive evolution.

## Methods

### RNA extraction, library generation and sequencing

Plants were grown in climatically controlled greenhouse conditions (12 h daylight, 25/20 °C day/night temperature) at the Arthur Willis Environmental Centre (AWEC) at The University of Sheffield. *Alloteropsis semialata* (AUS1 [Australian C_4_], RSA5-03 [South African C_3_], TAN1-04B [Tanzanian C_3_+C_4_] and ZAM1505 [Zambian C_4_]) and *Alloteropsis angusta* (UGA4) accessions were sampled from a living collection maintained at AWEC. *Themeda triandra* samples were taken from a Filipino accession^50^ (TtPh16-4), and *Setaria italica* was grown from seed (Herbiseed, UK). These species were chosen as *T. triandra* and *Setaria* sp. are known donors for some of the laterally acquired genes^32^. Samples from leaf tip, leaf base, leaf sheath, and root were taken on the same day, flash frozen in liquid nitrogen and kept at -80°C. Total RNA was extracted using a Qiagen RNeasy plant kit (Qiagen, Hilden, Germany) with an on-column DNA digestion step (RNase-Free DNase Set; Qiagen). RNA sequencing was performed in two batches. Libraries for the first 28 samples were generated using the TruSeq RNA Library Preparation Kit v2 (Illumina, San Diego, CA, USA), and 100 bp paired-end sequenced on two Illumina HiSeq 2500 flow cells (pooled with 20 samples from an unrelated project) at the Sheffield Diagnostic Genetics Service. For the remaining 46 samples, libraries were prepared and sequenced by Novogene, generating ∼5 Gb of 150 bp paired-end reads per sample using the Illumina NovaSeq 6000 s4 platform.

### Quantifying gene expression

Read quality was assessed before and after trimming with Fastqc v.0.11.8^51^ and Multiqc v.1.14^52^. RNA-seq data was cleaned using Trimmomatic v.0.38^53^ to remove adaptor contamination, low quality or ambiguous bases from both ends of each read (Phred quality score < 3), trim any remaining low quality bases (4 bp sliding window with mean Phred quality score <30), and finally remove short reads (<50 bp). The *Alloteropsis* RNA-seq data was mapped to reference genomes previously assembled for the same individual: AUS1 (GCA_004135705.1), RSA5-03 (GCA_036972165.1), TAN1-04B (GCA_036785585.1), ZAM1505 (GCA_036785565.1), UGA4 (GCA_037127165.1). *S. italica* was mapped to the Setaria_italica_v2.0 reference (GCA_000263155.2). *T. triandra* was mapped to *Sorghum bicolor* (Sorghum_bicolor_NCBIv3 GCA_000003195.3), a closely related grass from the same subfamily (Andropogoneae), as the only reference for *T. triandra* is fragmented (N50 = 13.4 kb), incomplete (81.6% BUSCO complete) and unannotated^50^. Paired-end reads were quasi-mapped onto their respective reference genome and transcript expression counts quantified using Salmon v.1.4.0^54^. Transcript counts were converted to gene level transcripts per million (TPM) using the R package tximport v.1.3.9^55^.

All sequencing datasets were also mapped to the *A. semialata* AUS1 reference genome for batch effect analysis. Principal component analysis (PCA) was undertaken using the package DESeq2 v.1.40.0^56^ to compare variability in expression profiles between samples. The resulting PCA did not highlight an effect of sequencing batch, but showed tissue-specific clustering, particularly separating leaf (sheath, base and tip) samples from root samples (SI Appendix, Figure S4).

### Identifying vertical and donor orthologs

This study uses laterally acquired genes identified by Raimondeau et al^35^. We restricted our analysis to only those present in the genome annotation (i.e. not those additionally identified using Blast^57^ by Raimondeau et al^35^), or those that were 100% identical that could cause problems quantifying their expression (one identical copy was removed). We revisited the trees generated by Raimondeau et al.^35^ to classify each laterally acquired gene into three groups: [1] vertically inherited homolog and donor xenolog present; [2] vertically inherited homolog present, donor xenolog absent; and [3] both vertically inherited homolog and donor xenolog absent. For donors, we only considered transfers from Cenchrinae and Andropogoneae, with either *S. italica* or *S. bicolor* sequences present, respectively. Corresponding vertically inherited homologs had to belong to the same accession.

### Laterally acquired gene expression analysis

For the gene expression analyses presented in the main text, we summed the TPM values of recent paralogs (i.e. sequences that formed a monophyletic *Alloteropsis* clade in the gene tree). We then used this to calculate log-TPM differences between laterally acquired genes, vertically inherited homologs and donor xenologs (Table 1). The log-TPM differences were calculated independently for each RNA-seq library, and we combined all tissues into each analysis. To determine if summing paralog expression values influenced our results, we repeated all analyses using: [1] only the most highly expressed paralog, and [2] only considering transfers without paralogs, i.e. neither the laterally acquired gene, vertical homolog or donor xenolog were duplicated in the gene tree (Table S2). Given the separation of root versus leaf tissues on principal component 1 (SI Appendix, Figure S4), we also repeated all analyses excluding roots (Table S2). In the main text we only refer to the results using summed TPM values across all tissue samples, but all the additional analyses are presented in Table S7, with only the coding sequence methylation analysis having predominantly conflicting results. All statistical analyses were conducted in R (version 4.4.1) using the dplyr^58^ and broom^59^ packages for data manipulation. Linear mixed-effects models (LMM) were fitted using the lme4 package in R^60^. To test for a significant difference in expression between laterally acquired genes and vertically inherited homologs we fitted a LMM with the following structure:

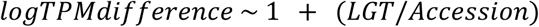

Where accession and LGT orthology were included as random effects to account for non-independence of our data. Model fitting was performed using restricted maximum likelihood (REML), and statistical significance was assessed using the lmerTest package^61^. To test for a difference in expression between laterally acquired genes and their xenologs in the donor, another LMM was fitted with the following structure:

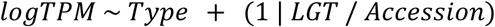

Where *Type* (laterally acquired or donor xenolog) is included as a fixed effect. The non-independence of the data is again accounted for as above.

To classify each laterally acquired gene as either ‘putatively functional’ or ‘degenerating’ we conducted two pairwise comparisons: [1] paired Wilcoxon signed-rank test to compare logCount between laterally acquired gene and vertically inherited homolog and [2] unpaired Wilcoxon rank-sum test between laterally acquired gene and donor xenolog. Multiple testing corrections were performed using the BH method^62^. Laterally acquired genes with significantly lower expression compared to both vertical homologs and donor xenologs were categorised as ‘degenerating’.

### Truncation analysis

To determine if the laterally acquired or vertically inherited orthologs were potentially non-functional, we compared their coding sequences in *Alloteropsis* to that of five other model grasses: *Brachypodium distachyon* (GCA_000005505.4)*, Oryza sativa* (GCA_001433935.1)*, Setaria italica* (GCA_000263155.2)*, Sorghum bicolor* (GCA_000003195.3)*, Zea mays* (GCA_902167145.1). Blastn^57^ was used to identify the top-hit match in each of the model grasses, before the full length sequences were aligned using the Geneious Prime 2024.0 (https://www.geneious.com) multiple sequence aligner. Each alignment was then manually inspected, and the *Alloteropsis* sequences classified as truncated if: [1] the *Alloteropsis* sequence was < 70% of the length of the other sequences; [2] no start codon at the beginning of the *Alloteropsis* sequence (present in model grasses); [3] no stop codon at the end of the *Alloteropsis* sequence (present in model grasses).

A GLMM was used to examine the relationship between truncation and whether the gene is laterally acquired or vertically inherited. The model was specified as follows:

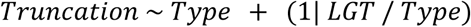

Where *Truncation* represents a binary option, and to account for the hierarchical structure of the data, *LGT* (parology) and *Type* (laterally acquired or vertically inherited) were included as random intercepts. The model was fitted using the glmer() function from the lme4 package in R^60^, with a binomial family and logit link function. A similar GLMM was additionally used to evaluate the effect of truncation status on laterally acquired gene expression category (degenerating vs putatively stable), specified as:

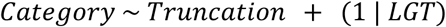

### Methylation

DNA was extracted from the four *A. semialata* accessions and sent to Novogene for whole genome bisulfite library preparation and sequencing. For each library, ∼35 Gb of data was generated (150 bp paired-end reads) using the Illumina NovaSeq 6000 S4 platform (coverage 33.2x - 50.0x ;Table S8). The bisulfite reads were trimmed using TrimGalore v0.6.10, a wrapper script that automates trimming (adapter and low quality bases), as well as the removal of biased methylation positions using Cutadapt v2.6^63^ and FastQC. The methylation analysis was performed using Bismark v0.24.2^64^. First, the reference genomes were indexed using the bismark_genome_preparation script, transforming the assembly into a bisulfite-coverted version (C-to-T and G-to-A converted). Then, the Bismark aligner performed read mapping for each individual to its respective bisulfite-treated reference genome, using Bowtie2 v2.4.1^65^. Methylation data extraction was carried out using Bismark’s methylation extractor tool, with the following options: --no_overlap --comprehensive --bedGraph. Methylated cytosine site information was extracted from the bedGraph file and filtered to remove cytosines with fewer than 10 or more than 50 reads. These data were further filtered to identify methylated cytosine sites within the whole gene body, the coding sequence, and 1 kb up- and down-stream from the gene. The proportion of methylated cytosines was calculated based on the total number of cytosines with aligned reads in the region and the number of cytosines where at least 50% of reads were methylated. To compare the proportion of methylated sites between laterally acquired and vertically inherited homologs, and between degenerating and putatively functional laterally acquired genes (based on gene expression only), we fitted LMMs using the lme4 package in R^60^. Accession and LGT orthology were included as random effects to account for non-independence of our data

### Selective constraint analysis

For the 116 laterally acquired genes that have a vertically inherited homolog present, we inferred the selection pressure they have been evolving under. A single representative from all *Alloteropsis* accessions was used as a reference, selected based on overall gene length and with a preference for selecting the same accession for both homologs. Pairwise dN, dS and dN/dS (ω) were calculated using the Yang and Nielsen^66^ method implemented in yn00, part of the PAML v.4.10.7 package^38^. To detect signatures of adaptive or purifying selection, we further applied both branch and branch-site models in CodeML, following Yang et al.^16^ protocol. We used the sequence alignments and phylogenetic gene trees from Raimondeau et al.^35^, filtered for full length sequences. For direct comparison between the lateral and vertical homolog, ingroup taxa were removed to make the two homologs sister to each other in the tree. We then independently set the branches leading to each of these as foreground branches. The inferred ω reflects the cumulative selection pressure acting on each gene since the point of divergence. For the laterally acquired genes, this includes both the evolutionary history within the donor lineage, and the time since integration into the recipient genome. To test for significant shifts in ω, a likelihood ratio test was performed against the null model (single ω across all branches). For branches under significant relaxed selection (i.e. significantly higher ω) we subsequently ran branch-site models (parameters: model = 2, NSsites = 2, fix_omega = 0) to identify codons under positive selection compared to the null model (fix_omega = 1).

### Molecular dating

The time since transfer for each laterally acquired gene was extracted from Raimondeau et al.^35^ to test if this is correlated with expression level or whether the gene is functional. To do this, we fitted a LMM using the lmer function from the lme4 package in R^60^. Random intercepts were included for laterally acquired genes and nested accession to account for hierarchical structure and repeated measures within laterally acquired genes and accessions. To assess the relationship between time since acquisition and our expression categories, we fitted a GLMM using the glmer function from the lme4 package in R^60^. Again, a random intercept was included for laterally acquired genes to account for non-independence among LGTs within the same laterally acquired gene. Both models were assessed for model significance using likelihood ratio tests. The molecular dating analyses were also performed at the individual level.

## Acknowledgements

This work was funded by a Natural Environment Research Council (NERC) grant (NE/V000012/1). CFC is supported by a University of Sheffield PhD scholarship, GS is supported by the MAPAS project (ERC grant agreement no 947921), P-AC is supported by a Royal Society University Research Fellowship (URF\R\180022), LP is supported by a NERC Standard Grant (NE/V000012/1), LTD is supported by a NERC Independent Research Fellowship (NE/T011025/1).

## Author contributions

CFC, P-AC, LP and LTD designed the study. SGSH and PR sampled plants for RNA work. EB, LP, GS and PR performed RNA lab work. LTD generated bisulfite data. CFC analysed the data with the help of BTA, LP and LTD. All authors helped interpret the results. CFC, LP and LTD wrote the manuscript with the help of all authors.

## Data availability

All raw RNA and bisulfite sequencing data have been deposited with NCBI under Bioproject PRJNA1299365. All scripts used in this study are available on GitHub: https://github.com/Sheffield-Plant-Evolutionary-Genomics/.

## Supplementary Information for

### Supplementary Figures

**Supplementary Figure 1.**
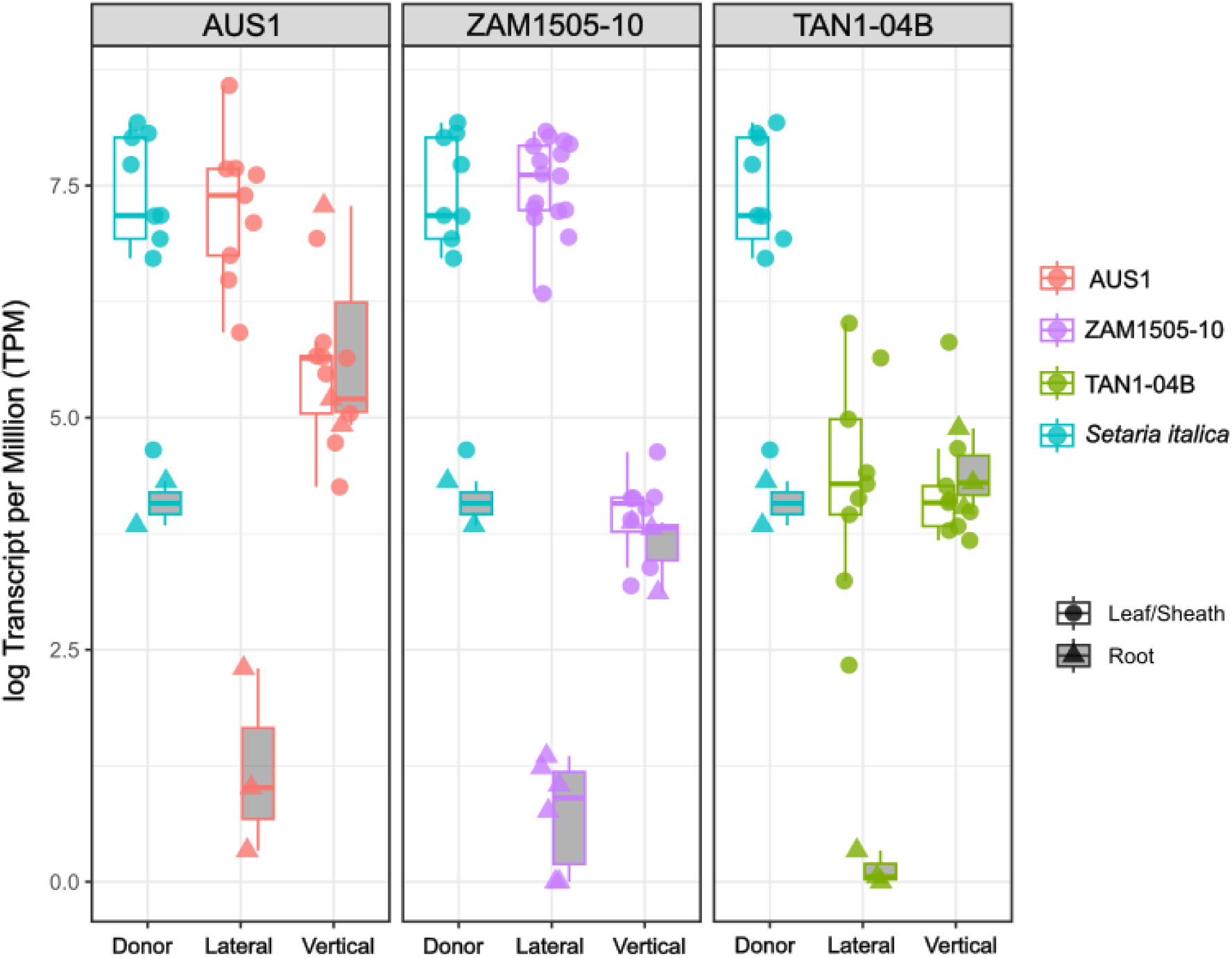
Expression patterns of LGT-019, phosphoenolpyruvate carboxykinase (PCK). Boxplots show median and interquartile range

**Supplementary Figure 2:**
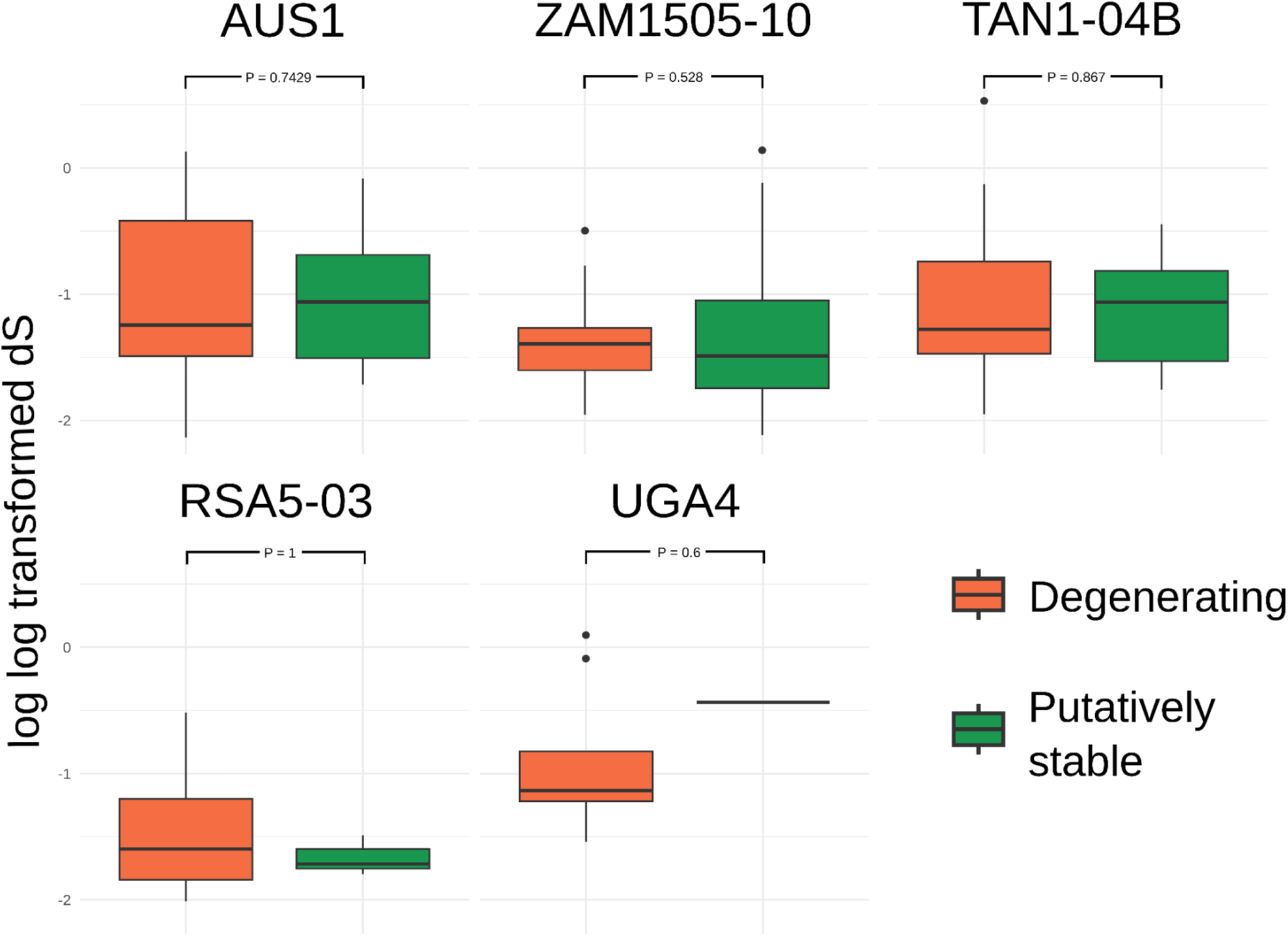
Genetic divergence between corresponding vertically inherited and donor proxy genes for laterally acquired genes classified as ‘Degenerating’ and ‘Putatively stable’. dS indicates synonymous substitutions between the gene pairs; this value is twice log transformed to create a normal distribution of the data.

**Supplementary Figure 3.**
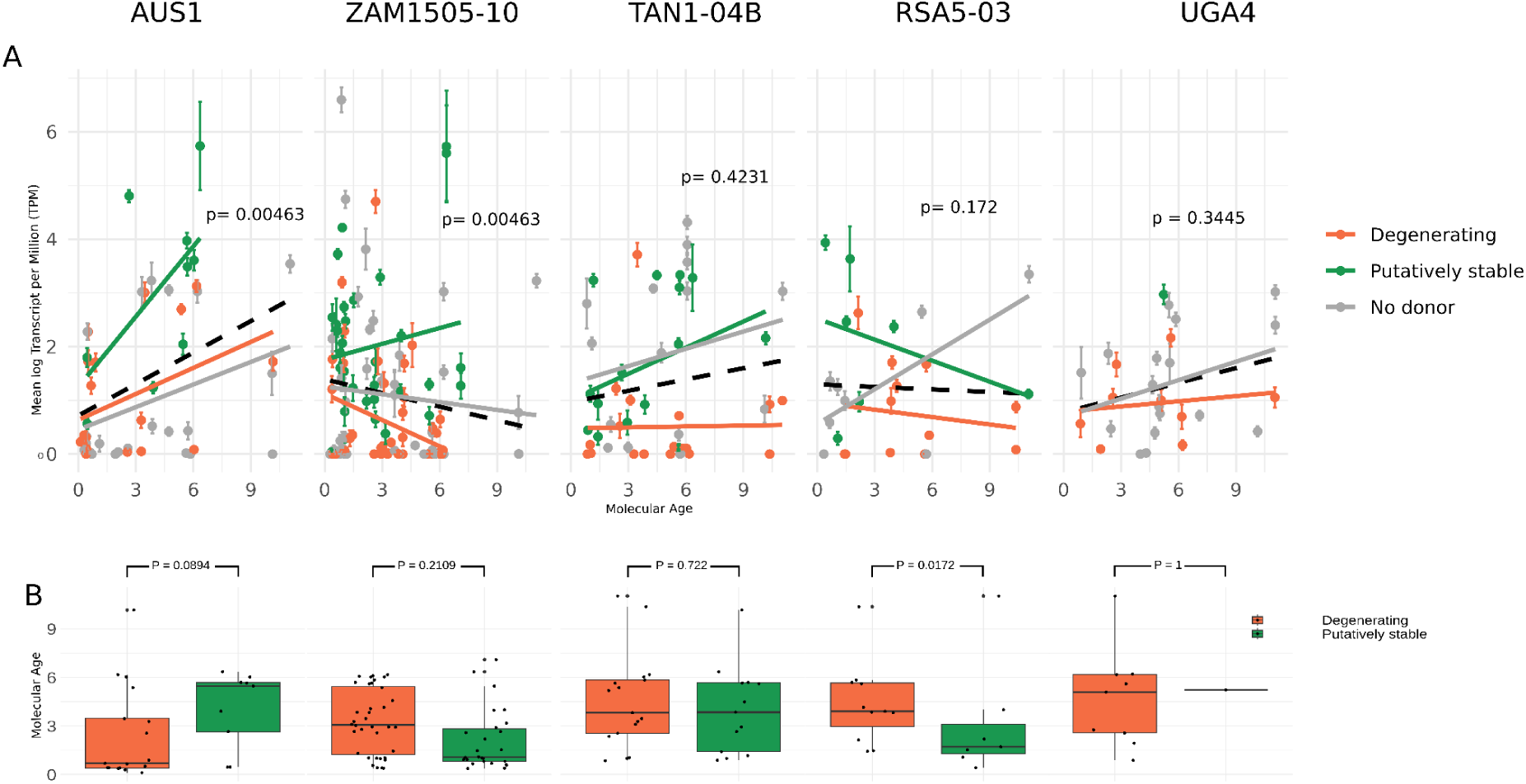
Time since acquisition and expression patterns of laterally acquired genes. A) Relationship between age and expression. Error bars indicate variation across replicates per laterally acquired gene. Black dashed line indicates the linear relationship between age and TPM, with p values from linear mixed effects models included. Coloured lines indicate trends for laterally acquired genes classified as ‘degenerating’ or ‘putatively stable’. No donor indicates laterally acquired genes where a donor proxy gene was not found in the gene phylogeny. B) Boxplots showing the range of molecular age across laterally acquired genes within expression categories. P values from Wilcoxon Signed Rank tests are included above boxplots.

**Supplementary Figure 4:**
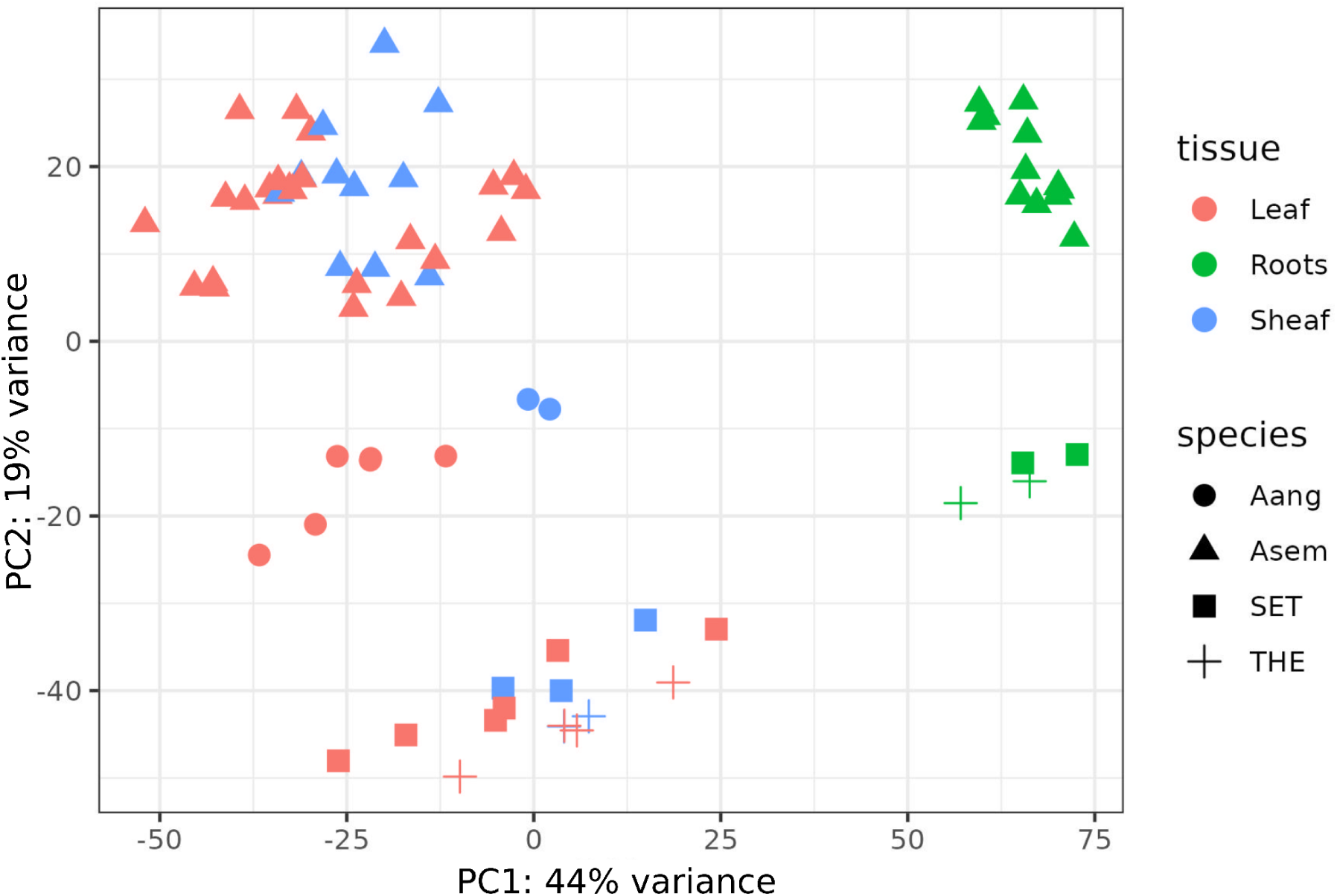
Principal component analysis (PCA) of RNA-seq data demonstrates tissue effect between the leaf and sheath tissue types (red and blue respectively) and the roots (green). Aang = *Alloteropsis angusta*, Asem = *Alloteropsis semialata*, SET = *Setaria italica*, THE = *Themeda triandra*.

